# Classification of bioactive peptides: a comparative analysis of models and encodings

**DOI:** 10.1101/2023.10.04.560809

**Authors:** Edoardo Bizzotto, Guido Zampieri, Laura Treu, Pasquale Filannino, Raffaella Di Cagno, Stefano Campanaro

**Affiliations:** Department of Biology, University of Padua, via U. Bassi 58/b, 35131 Padova, Italy; Department of Soil, Plant and Food Science, University of Bari Aldo Moro, Via G. Amendola 165/a, 70126 Bari, Italy; Faculty of Agricultural, Environmental and Food Sciences, Free University of Bolzano, piazza Universita, 5, 39100, Bolzano, Italy

**Keywords:** bioactive peptide, functional classification, machine learning, sequence encoding, systematic evaluation

## Abstract

Bioactive peptides are short amino acid chains possessing biological activity and exerting specific physiological effects relevant to human health, which are increasingly produced through fermentation due to their therapeutic roles. One of the main open problems related to biopeptides remains the determination of their functional potential, which still mainly relies on time-consuming in vivo tests. While bioinformatic tools for the identification of bioactive peptides are available, they are focused on specific functional classes and have not been systematically tested on realistic settings. To tackle this problem, bioactive peptide sequences and functions were collected from a variety of databases to generate a comprehensive collection of bioactive peptides from microbial fermentation. This collection was organized into nine functional classes including some previously studied and some newly defined such as immunomodulatory, opioid and cardiovascular peptides. Upon assessing their native sequence properties, four alternative encoding methods were tested in combination with a multitude of machine learning algorithms, from basic classifiers like logistic regression to advanced algorithms like BERT. By testing a total set of 171 models, it was found that, while some functions are intrinsically easier to detect, no single combination of classifiers and encoders worked universally well for all the classes. For this reason, we unified all the best individual models for each class and generated CICERON (Classification of bIoaCtive pEptides fRom micrObial fermeNtation), a classification tool for the functional classification of peptides. State-of-the-art classifiers were found to underperform on our benchmark dataset compared to the models included in CICERON. Altogether, our work provides a tool for real-world peptide classification and can serve as a benchmark for future model development.

## Introduction

Bioactive peptides (BPs) are short chains of 2 to 50 amino acids with a molecular weight of less than 10 kDa exerting biological effects on unicellular and multicellular organisms [1,2]. Their biological properties make them useful as potential candidates for the treatment of pathologies where conventional medication is ineffective or not advised [3,4]. BPs can have several beneficial functions such as anti-inflammatory, antihypertensive and antidiabetic [1]. Additionally, they can possess antibacterial, antiviral, and antifungal properties that are comparable to, or even surpass, those of antibiotics in terms of efficacy [5,6]. BIOPEP-UWM, a database focused on cataloging BPs, currently lists 62 different functional classes, which can include a single peptide up to hundreds of individual molecules [7]. BPs can derive from proteolysis where the primary sources are proteins derived from plants, animals, and even marine species [8]. The most extensively studied sources of precursor proteins are those used for human nutrition, including milk and other dairy products, soybean and green leafy vegetables [9,10]. By-products derived from agrifood processes are being considered as an additional source of BPs; their use is of particular interest because extraction of high-value biomolecules can open up a new way for the recycling of waste products, such as animal skin and feathers, or fruit peel [11,12]. There are three main methods to obtain bioactive peptides: chemical and physical hydrolysis, microbial fermentation and enzymatic hydrolysis [13]. In the former, proteins are placed in acidic or basic environments at different temperatures or are subjected to microwave irradiation or ultrasonic treatment to break down amino acid chains [14]. In the latter, single or multiple peptidases from animal or plant sources are added to the substrate to obtain BPs [15,16]. In substrates fermented by bacteria and fungi, microbial peptidases hydrolyze amino acid chains [17,18]. After converting the protein matrix into smaller peptides, the sequence of the BPs can be determined through chromatographic purification and mass spectrometry. Subsequently, to verify and measure the functional activity of BPs, a series of steps, including purification and in vivo testing, must be performed. However, these procedures are time-consuming and incur significant monetary costs.

Given the importance of BPs, there have been several attempts to create in-silico approaches to perform a preliminary assignment of the potential functional properties and facilitate the subsequent discovery and testing process in vivo [19–24]. These methods rely on several databases where peptides from various experiments have been collected and classified according to the BPs functional classes. Using the sequence properties of the peptides, such as amino acid composition, or the presence of sequence patterns of interest, peptides can be assigned to a functional class depending on the type of classifier used. Tools include similarity-based classification using available sequences from databases [7,25] and prediction of physicochemical properties [26]. There have also been several attempts to use machine learning techniques to aid in the detection of BPs and their functional classification [19–24]. The proposed methods used Logistic Regression [27], Support Vector Machines [28] and Random Forests [29] to predict the BPs functional role. In recent years, neural networks [22,30,31] and algorithms based on natural language processing (NLP) have also been employed for the same purpose. These studies tend to focus solely on a single method for classification and in most cases they do not compare the performance of the model with other machine learning algorithms. While the intrinsic characteristics of bioactive peptides require the use of different techniques for classification, there exist no universally accepted benchmark settings for testing this type of problem. Moreover, in order to use these different classification methods, peptide sequences need to be transformed into appropriate signals that can be processed by the algorithm of choice. Various encoding techniques have been developed to take into account the different properties of proteins [32,33]. For instance, amino acids can be encoded by assigning them a number based on the order of the 20 conventional amino acids, or by calculating their frequency in one sequence. Other methods use the physicochemical parameters or secondary structures of BPs. While these techniques have been proven to be useful in transferring information from the sequence to the machine learning algorithm, they sometimes fail to convey important details, such as physical or chemical properties or relationships occurring among amino acids. This leads to predictors that are highly condition-specific and that cannot be generalized to problems other than the ones they were developed for [34].

The aim of this study is the generation of a set of classifiers for the identification of peptide functions, with a particular focus on BPs derived from microbial fermentation. There are several reasons why microbial fermentation is preferable to enzymatic or physicochemical methods of extraction. First and foremost, utilizing microbial species is far cheaper than traditional methods and does not require solvents or exogenous enzymes, making it a green and sustainable alternative. Moreover, the high number of species and the variability in the enzyme portfolio produce a higher number of bioactive peptides of different sizes. As far as the authors are aware, there are no studies focusing specifically on BPs from microbial sources with the exception of the formation of specific databases focused on fermented food peptides. For this reason, we propose CICERON, a tool to classify the functions of BPs specifically derived from microbial fermentation.

Different machine learning and encoding methods were evaluated for a total of 171 distinct classifiers; this approach allowed to highlight the differences in the techniques employed and their performance in evaluating the individual classes. The various encodings and machine learning techniques were tested to provide a benchmark for future studies focusing on peptides derived from microbial fermentation. For each functional class of BP, the most accurate predictor has been selected to suit the intrinsic characteristics of each function. A Python script was generated to identify the functions of BPs. The final result, CICERON, consists of nine different binary classifiers capable of identifying the products of microbial fermentation-derived BPs. The database of microbial peptides used in this study and the best model for each functional class can be found at https://github.com/BizzoTL/CICERON/

## Methods

### Dataset preparation

Peptide data from BIOPEP-UWM [7], Milk Bioactive Peptide Database [35], BioPepDB [36] and FermFoodDB [25] were downloaded and merged into a unified database. Data collected from these databases are represented only by food proteins derived from microbial fermentation. BPs that were present more than once due to overlaps in the online databases were clustered together to remove redundancy. Peptides with identical sequences but different functional class assignments were removed to avoid introducing potential biases in the classifier’s training. Additionally, sequences that had more than 90% similarity were also clustered together if the peptide belonged to the same functional class, otherwise, they were excluded from the analysis. Peptides longer than 100 amino acids, shorter than 3 and those containing unconventional amino acids or symbols were also removed. Functional classes were homogenized by grouping together those with the same or overlapping biological functions, as follows: “antihypertensive”, “ACE-inhibitory” and “Renin-inhibitory” as Antihypertensive; “DPP-IV inhibitors” and “alpha-glucosidase inhibitors” as Antidiabetic; “antimicrobial”, “antifungal”, “antibacterial” and “anticancer” as Antimicrobial; “antithrombotic”, “CaMKII Inhibitor” as Cardiovascular with positive effects on vascular circulation; “Antiamnestic”, “anxiolytic-like”, “AChE inhibitors”, “PEP-inhibitory” and “neuropeptides” as Neuropeptides. Among the functional classes mentioned above, celiac disease is the only one that determines a toxicity-inducing effect, contrary to all the others that have a positive effect on human health, due to their allergenic properties. Finally, classes with less than 100 peptides were removed.

### Functional class characterization

Amino acid, dipeptide and tripeptide frequencies were calculated to infer possible relationships between functional classes and amino acid composition. MERCI software [37] was used to discover sequence motifs unique for each class. A binomial test and false discovery rate correction were carried out to infer the statistical significance of the results, in order to ascertain whether the most frequent dipeptide or tripeptide sequences were more prevalent in one class compared to others. For each sequence of interest, the probability of finding that specific motif was calculated across all samples and compared to the distribution of the sequence within the class of interest.

### Encoding methods

BPs were encoded into appropriate inputs for different machine-learning algorithms (Figure 1). Four different encoding methods were devised for Support Vector Machine (SVM), Random Forest (RF), K-Nearest Neighbor (KNN) and Logistic Regression (LR) models. In the “sparse” encoding method, each peptide was transformed into a vector of length 100, representing the maximum possible sequence length in the database. Using a list of the 20 conventional amino acids and an additional element representing empty positions, each amino acid in the sequence was encoded as a 21-element vector. The position of the amino acid in the list was assigned a value of “1,” while the remaining 20 elements were set to “0.” Thus, for each position in the vector of length 100, there was a corresponding vector of length 21. The “dense” encoding method also utilized a vector of length 100. However, instead of one-hot encoding for each position, the indices from 1 to 20 representing the 20 conventional amino acids were used, along with an element indicating an empty position. The third encoding method, “BLOMAP” [38], used the BLOSUM62 substitution matrix to encode each amino acid as the vector of the substitution probability with other amino acids. The additional character indicating a missing amino acid was given a -100% probability of substitution with the other 20 amino acids. The last encoding method, “thremers”, involved using a sliding window of 3 amino acids to count all the 3-mers in the peptide. These counts were then stored in a dictionary encompassing all possible combinations of the 20 common amino acids. The Neural Network (NN) models used “sparse” and “threemers” encodings to map the input for the first convolutional layer. The encoding for the BERT protein learning model, ProtBERT [39], consisted in separating each sequence into single amino acids and treating them as a single element, as follows. The maximum length of the encoded vector was set to 102 elements. The first element in the vector was a special character that indicates the start of the peptide, followed by the tokenized amino acids, and ending with the special character indicating the end of the sequence. Any remaining positions were padded to ensure uniform vector lengths of 102. The entire peptide sequence is treated as an entire phrase in the language model. All the BPs were encoded with the methods described above and a t-SNE [40] representation was used to infer information on the sequence distribution of functional groups (Supplementary Figure 1).

**Figure 1.**
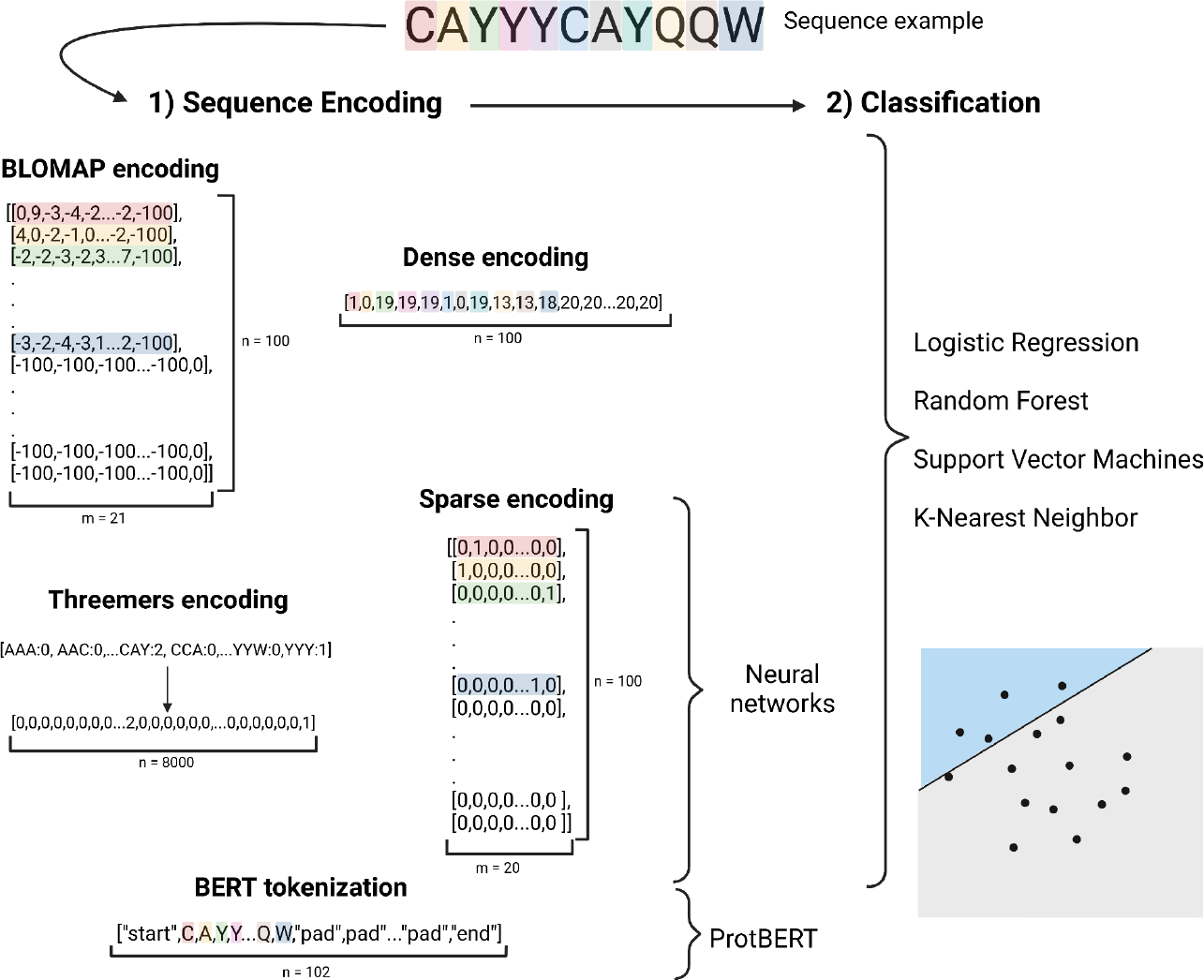
Example of classification of a bioactive peptide sequence. 1) The sequence was first transformed into the appropriate vectors according to the encoding. 2) The encoded sequence was then used in the classification by different algorithms.

### Classifiers

For every functional prediction, the samples belonging to the specific functional group under investigation were labeled as “0”, while the remaining samples were labeled as “1”. The data was encoded using the four encoding methods described previously. LR, SVM, RF and KNN were implemented in the Scikit-learn Python Package version 1.1.3 [41]. Convolutional NN were implemented using the Keras framework version 2.11.0 [42]. The architecture consists of two convolutional layers, each followed by two Max pooling operations. This was then followed by a Dropout layer and a flattening operation. Two Dense layers were added, with the last one being the output layer. For both traditional classifiers and NN the dataset was split into training, validation, and test sets in a ratio of 70:20:10. The number of epochs was set to 50, and the batch size was set to 20. The encoding methods tested for the CNN models were “sparse” and “threemers”. The protein learning model, based on the ProtBERT pre-trained model, was implemented using the HugginFace Transformers framework version 4.26.0 [43]. Each binary classifier was trained using an 80:20 training test split for 20 epochs. After each epoch, the trained classifier was tested on the validation set and the resulting metrics were recorded. Among all the machine learning methods tested, the model with the best MCC obtained from the test set was selected as the representative for each class.

### Model evaluation and parameter optimization

Due to the significant imbalance in the number of BPs present in the different classes, accuracy alone cannot adequately describe the performance of the models. The metrics used in the classification were therefore the Area Under the Receiver Operating Characteristic curve (AUROC) and Matthew’s Correlation Coefficient (MCC). Each of the functional classes in the database has been predicted using a binary classifier for each machine learning algorithm previously described. The following parameters were selected in order to optimize the performance of the model: “C”, “penalty”, “solver” for LR; “criterion”, “max_features”, “n_estimators” for RF; “algorithm”, “leaf_size”, “n_neighbors”, “weights” for KNN; “C”, “class_weight”, “gamma”, “kernel” for SVM. A grid search was performed to select the best parameters for each model, followed by cross-validation, using the GridSearchCV module from scikit-learn. The best parameters obtained for each model are described in Supplementary Table 1. For NN models, a genetic algorithm was developed using the DEAP Python library [44]. The weights of the positive and negative classes, learning rates, and kernel size were encoded as a vector and used as individuals in the genetic algorithm. The MCC obtained from the test set served as the fitness value for the individuals. After each generation, the top 5 individuals with the highest fitness were selected as parents for producing offspring in the next generation. A randomly selected individual from the pool of the best individuals had a 25% chance of changing each hyperparameter using a random value. The allowed intervals for the class weights ranged from 1 to 10 (float values), for the learning rates it ranged from 0.0001 to 0.1, and for the kernel size it ranged from 5 to 500 (integer values). For the first 10 generations, the number of offspring was set to 15, while the remaining 5 individuals were randomly generated within the set of possible intervals. After the 10^th^ generation, all 20 individuals were generated as offspring from the best individuals of the previous generations. In order to select the best hyperparameters for the singular classes, the genetic algorithm consisted of 20 generations of 20 individuals each for every functional class. The vector used to obtain the best neural network model for each class is described in Supplementary Table 1. The metrics obtained from the best individual of the 20 generations were further evaluated using 5-fold cross-validation to obtain the final metric values.

### CICERON implementation

CICERON consists of a Python script that takes one or more FASTA files as input and returns the functional prediction for every peptide reported in the file. The first step consists of checking that the input sequence is present in the database of bioactive peptides and if the match is identical the function is reported in the final table. The second step consists of the search for motifs: if the sequence contains one of the class-specific motifs previously found using MERCI, the functional class is reported. In each step, if multiple associations are confirmed, they are all included. Finally, the best classifier model for each functional class predicts the probability of the peptide belonging to that group or not. The results are collected in a tab-separated file for each file in input.

## Results and discussion

### Exploration of the dataset and functional class properties

In our starting dataset, a total of 13,123 peptides were obtained from different online databases, which were divided into 56 functional groups. After filtering the sequences, and merging functionally overlapping or related classes, the final database consisted of 3990 BPs divided into nine different functional groups (Table 1). Such classes present substantial differences both in terms of number of peptides and sequence characteristics. The number of BPs per class ranges between 107 for “immunomodulatory” to 1386 for “antihypertensive”. The difference in the number and the length of BPs for each class reflects the difficulty in performing certain essays for the identification of functions in vivo and the higher interest in certain functions over others. It is also possible that some biological functions can be performed only by a very limited number of AA sequences, thus intrinsically limiting the number of BP in the class. Moreover, after the production of the digested substrates, peptides are filtered from other molecules based on the molecular weight, prioritizing shorter peptides over longer ones and thus reducing the number of BPs that are usually longer, such as antimicrobial BPs. While the average peptide length is 9.5 amino acids, the length distribution is quite different between functional classes. For antidiabetic and immunomodulatory classes the peptide length is shorter than the global average, while for antimicrobial the peptides are significantly longer than the mean, more than double in comparison to other functional groups (Supplementary Figure 2). In the t-SNE representations (Supplementary Figure 1), the different classes are not clustered together, with the exception of celiac disease peptides in “sparse” and “threemers” encodings, where the majority of these samples are clustered in a unique group.

**Table 1.**
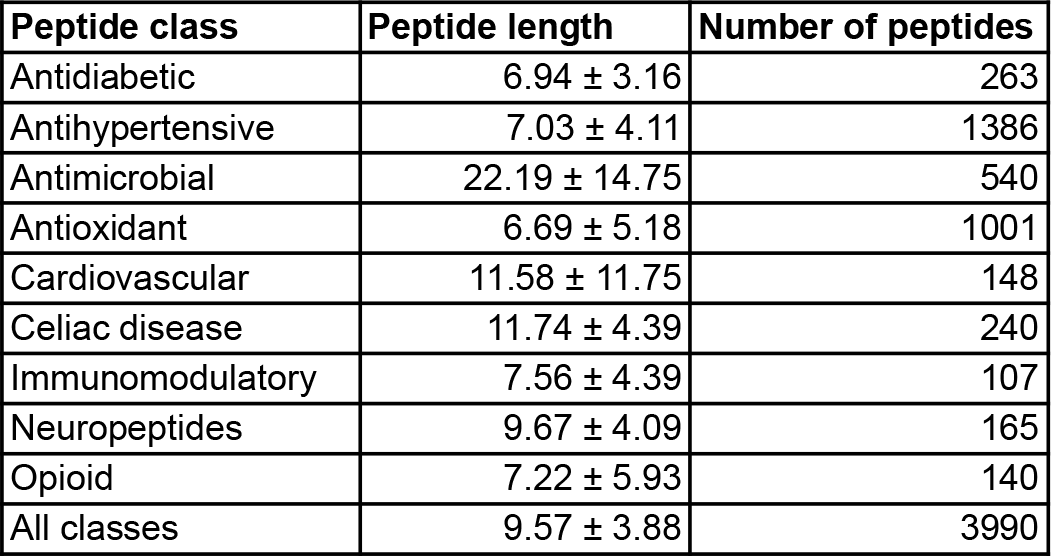
Average sequence length and number of peptides for each functional class.

To compare the amino acid usage in the functional classes, single amino acid, dipeptide, and tripeptide frequencies were plotted (Figure 2). The amino acid frequency plot in Figure 2A reveals that some BPs classes have distinct characteristics. For example, celiac disease BPs have the highest frequency of proline and glutamine, opioid peptides are enriched in tyrosine and glycine, while cardiovascular BPs have slightly higher frequencies of alanine and the highest frequency of the negatively charged amino acids aspartic acid and glutamic acid.

**Figure 2.**
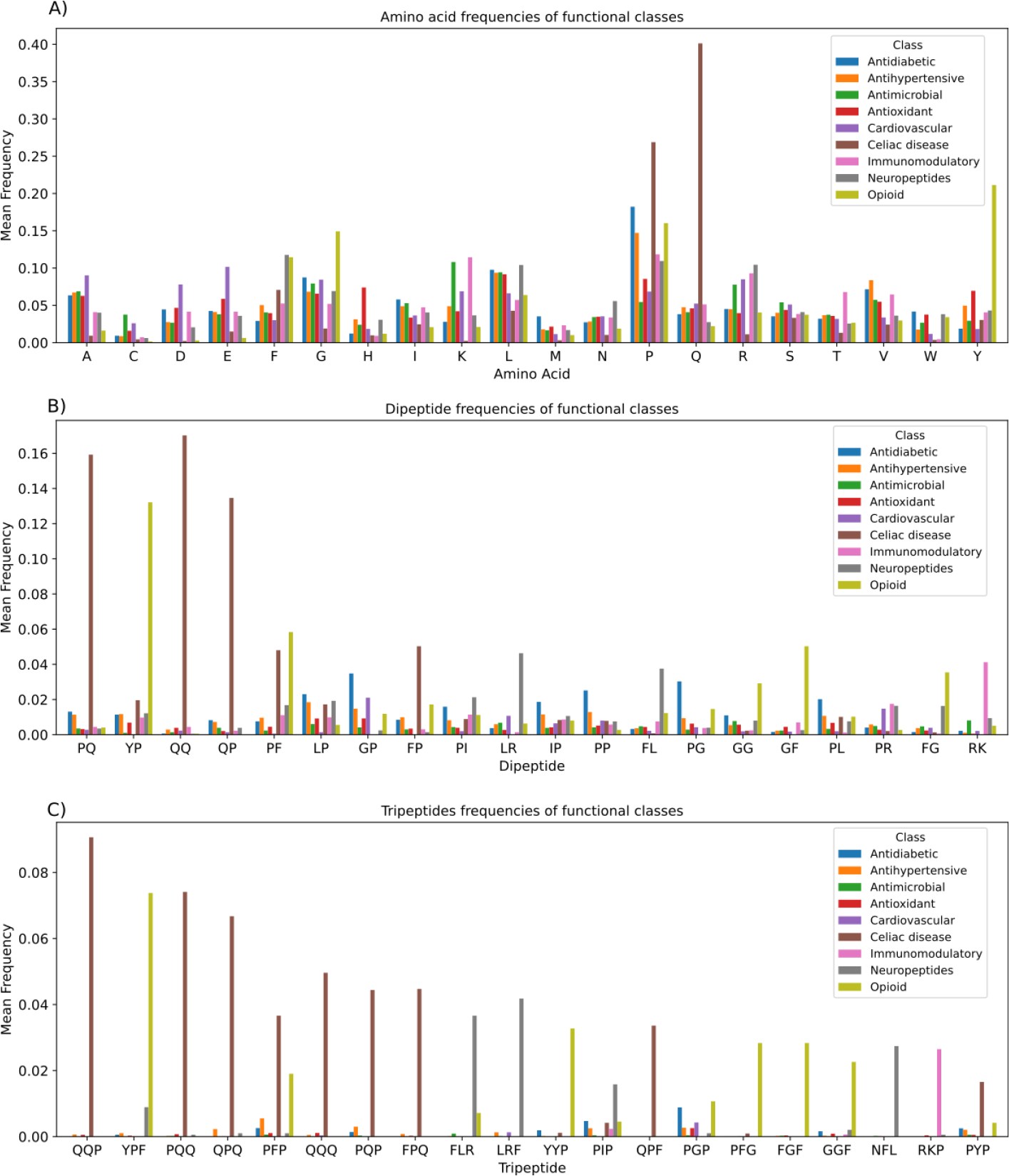
Amino acid composition of peptides for each functional class: A) Mean amino acid frequencies for single amino acids, B) Mean dipeptide frequencies of the 20 most abundant dipeptides, C) Mean tripeptide frequencies for the 20 most abundant tripeptides.

Distinct patterns are also visible in the frequency of dipeptides (Figure 2B), with high-frequency dipeptides mainly localized in celiac disease-associated and opioid BPs, immediately followed by antidiabetics. This trend is further confirmed by the tripeptide distribution (Figure 2C), where celiac disease and opioid peptides have 9 and 5 amino acid triplets, respectively, occurring at higher frequencies. The remaining most frequent tripeptides are from the neuropeptides class (FLR, LRF and NFL) and the immunomodulatory class (RKP). PIP and PGP are tripeptides that have a high frequency in multiple classes. A binomial test also identified that many other sequences were present with a higher frequency in certain classes, providing additional insights and potential patterns that can be exploited for classification (Supplementary Table 2).

Further analysis using the MERCI revealed unique motifs for some functional classes. For celiac disease, the QPF triplet was found 67 times in the 240 samples and 9 other motifs were found in at least 10% of the sequences. Glutamine and proline were prominent in these motifs, occurring at least once per motif. All the discovered motifs from MERCI are distinct for each class and are not found in samples that belong to a different function. Although unique motifs can be used to positively identify classes, they should not be solely relied upon for functional classification. In the work of Tomer et al. [45], QPQ triplet was described as highly conserved in celiac disease; however, in the database produced in this work, this motif was found in 21 non-celiac disease peptides. The presence of a motif alone does not unequivocally determine the functional class of a peptide. Another possible option is that these 21 peptides have multiple functions, including also celiac toxicity. The distribution of amino acid frequencies, along with specific substructures, can provide insights and aid in the identification and classification of bioactive peptides, but they should not be the sole method of peptides functional classification. All the motifs found for each class are reported in Supplementary Table 2.

### Benchmark model evaluation across functional classes

In order to establish a benchmark for the classification of BPs, combinations of several sequence encoding and machine learning techniques were tested across the nine functional classes (Figure 1). To address the class imbalance in the dataset, MCC was used as the evaluation metric for comparing the performance of different classifiers, as shown in Figure 3. The obtained AUROC was also reported in Supplementary Figure 3, and the performance metrics obtained during training and testing are reported in Supplementary Table 1.

**Figure 3.**
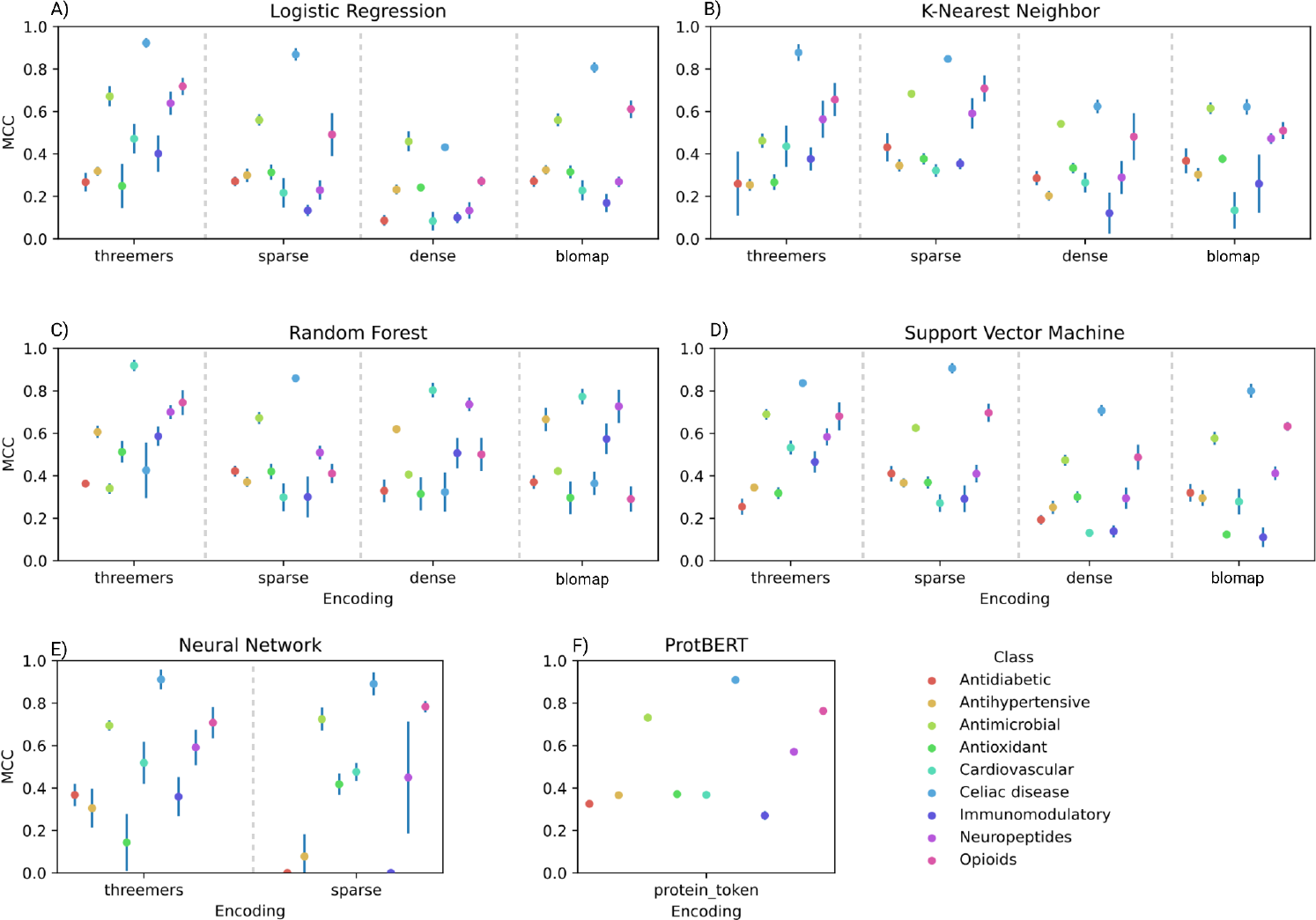
Matthews Correlation Coefficient value on the test set for every single classifier and encoding method. The dot size corresponds to a value of 0.03 MCC. Standard deviation values lower than 0.03 are not displayed. Each plot describes the performance of a different machine learning method: A) Logistic Regression, B) K-Nearest Neighbor, C) Random Forest, D) Support Vector Machine, E) Neural Network, F) ProtBERT.

The LR, RF, SVM and KNN algorithms were assessed in conjunction with “sparse”, “dense”, “BLOMAP” and “threemers” encoding methods. In general, each classifier exhibited varying levels of performance when coupled with the different encoding methods and no single encoding method outperformed the others, as evidenced by the findings presented in Figure 3. For example, the “threemers” encoding exhibited exceptional effectiveness in the classification of cardiovascular, immunomodulatory, and antioxidant classes, likely due to the presence of specific tripeptides that augmented the classification power. On the other hand, the “dense” and “BLOMAP” encodings yielded the most favorable MCC values for neuropeptides and antihypertensives, respectively. Despite such a heterogeneity, the “sparse” and “threemers” encoding methods yielded the highest number of classifiers with MCC values exceeding 0.5, and consequently, they were selected as the encoding methods for Neural Networks (NN) to reduce the time for computational training. Analogous trends were observed for NN models, with highly variable performance associated with individual feature types depending on the considered functional class. Overall, the variability in the optimal encoding method across all classes underscores the necessity for a robust benchmark encompassing a wide range of encoding strategies to ensure the comprehensive capture of relevant features in BPs classification.

When comparing classification model types, while no single machine learning method consistently excelled across all classes, RF demonstrated superior models for five out of nine classes, indicating its effectiveness in discovering relevant features within the dataset. KNN, on the other hand, displayed less favorable overall performance, with the exception of the antidiabetic class. However, the performance of this class, the one only under 0.5 MCC, is the lowest among all classes, indicating that the encoding methods are not sufficient to capture enough information to correctly classify these peptides. Notably, LR emerged as the optimal classification method for the celiac disease class, achieving the highest MCC value among all the classifiers at 0.923 with the “threemers” encoding. This excellent performance can be attributed to the high fraction of distinctive motifs in the class, which facilitates effective separation based on the presence of specific tripeptides, a characteristic more prevalent within this class (Figure 2 and Supplementary Table 2). Along with celiac disease BPs, opioids and antimicrobials display the highest scores on average, with MCC values of 0.783 and 0.724, respectively, achieved by NN in conjunction with the “sparse” feature encoding. This enhanced performance for antimicrobials was likely due to the significantly longer peptide sequences in antimicrobial peptides, averaging over 22 amino acids, four times the overall average length of all BPs. For the opioid class, the high performance is likely due to the presence of specific motifs, as it is the second class with the most frequent tripeptide motifs, after celiac disease, as seen in Figure 2. Moreover, we verified that the NLP-based ProtBERT model, pre-trained on a vast corpus of sequences, does not provide improved performances in the present tasks except for antimicrobial peptides, where ProtBERT exhibited a slight performance improvement over NN, achieving an MCC of 0.731. The length of antimicrobial peptides makes them more likely to be classified using the BERT-derived algorithm. However, it is worth noting that the use of the native ProtBERT tokenizer as an encoding method for such a model might have limited its performance in certain cases.

The systematic evaluation unveiled a high heterogeneity across functional classes, both in terms of distinctive sequence characteristics and model power. For this reason, no single classification algorithm or encoding method is capable of systematically outperforming others, highlighting the need for robust benchmark testing with multiple techniques to better elucidate the differences in class-specific features. Ultimately, the best-performing classification method and encoding for each class were selected as representatives for their respective classes and subsequently incorporated into the CICERON tool.

### CICERON comparison with state-of-the-art classifiers

To set a reference in BP classification research, CICERON was tested against state-of-the-art (SOTA) classifiers specific to the various peptide functional classes, which were selected upon careful literature inspection [45–50]. The main differences are summarized in Table 2. It is worth noting that certain functional classes, such as opioid, immunomodulatory, and cardiovascular peptides, could not be included in our comparative analysis due to the absence of available classifiers for these specific categories. Additionally, for the antihypertensive and neuropeptides classes, the tools described in relevant literature were not accessible for utilization. For the antimicrobial class, it is important to mention that the web server version of the tool allowed the submission of only one sequence at a time, while the source code version lacked comprehensive installation and usage guidance for the package. For the other three classes, namely antidiabetic, antioxidative, and celiac disease, the database of peptides generated in this study was used to compare the performance of these purpose-specific tools against CICERON.

**Table 2.** Comparison of state-of-the-art classifier and CICERON for each functional class. Opioid, immunomodulatory and cardiovascular classifiers were not included due to the absence of available classifiers. The characteristics that were considered are the following: model, encoding, type of tool and dataset used; MCC on the CICERON dataset, limitation of the tool.

| Functional Class | Tools | Model | Encoding | Type of tool | Training, validation and test dataset | MCC on CICERON dataset | Limitations | Ref |
| --- | --- | --- | --- | --- | --- | --- | --- | --- |
| Antidiabetic | AntiDMPpred | Random Forest | Multiple encodings were merged | Web tool | 236 antidiabetic and 236 non-antidiabetic peptides | 0.133 | Validated on peptides between 5 and 50 AA | [46] |
|  | CICERON | KNN | Sparse (One hot encoding) | Python package | 263 antidiabetic and 3727 non-antidiabetic peptides | 0.432 | / | This work |
| Celiac disease | CDpred | ExtraTree classifier | Amino Acid Composition (AAC). | Web tool | 503 celiac disease and 503 non-celiac disease peptides | 0.47 | / | [45] |
|  | CICERON | Logistic Regression | Threemers (Tripeptide composition) | Python package | 240 celiac disease and 3750 non-celiac disease peptides | 0.923 | / | This work |
| Antioxidant | AnOxPP | BiLSTM Neural Network | 22 Amino Acids Descriptors (AADs). | Web tool | 1060 antioxidant and 1060 non-antioxidant peptides | -0.005 | Applicable on peptide sequences between 2 and 19 AA | [47] |
|  | CICERON | Random Forest | Threemers (Tripeptide composition) | Python package | 1001 antioxidant and 2989 non-antioxidant peptides | 0.513 | / | This work |
| Neuropeptides | Target-ensC_NP | Ensemble of ETC, LGBM, SVM, XGB, and ADA | One-hot encoding of single AA | Python package | 2435 neuropeptides and 2435 non-neuropeptides | / | Tool not available for use | [48] |
|  | CICERON | Random Forest | Dense (One hot encoding) | Python package | 165 neuropeptides and 3825 non-neuropeptides | 0.736 | / | This work |
| Antihypertensive | Ensemble-A HTPpred | Ensemble of RF, SVM and XGB | 431 numerical features | Web tool | 913 antihypertensive and 913 non hypertensive peptides | / | Tool not available for use | [50] |
|  | CICERON | Random Forest | BLOMAP | Python package | 1386 antihypertensive and 2604 non hypertensive peptides | 0.665 | / | This work |
| Antimicrobial | Antimicrobial-peptide-generation | SeqGAN and BERT | BERT tokenization | Web tool and Python package | 4134 AMPs and 4134 non-AMPs | / | Validated on peptides between 11 and 30 AA; web tool only accepting one sequence at once; python package lacking a guide for utilization and requiring retraining the model | [49] |
|  | CICERON | ProtBERT | BERT tokenization | Python package | 540 AMPs and 3450 non-AMPs | 0.731 | / | This work |

Results reveal that CICERON outperformed SOTA models across all the three comparable classes. Specifically, CICERON exhibited MCC values that were consistently 30-50% higher than those of the benchmarked SOTA models. The notable difference observed in the antidiabetic classifier’s performance can be attributed to the relatively constrained dataset employed, characterized by a limited number of samples for both positive and negative classes, thereby restricting the model’s classification capabilities. In contrast, the lower MCC values observed for the celiac disease and antioxidative classes primarily stem from an increased number of false negatives in the predictions. It is worth mentioning that a portion of the positive samples for these classes was collected from the same databases utilized for data generation in this study. Consequently, it is the negative dataset that fundamentally contributes to the different performance of the tools.

Most of the SOTA class-specific classifiers are trained on a large dataset of positive examples and a random dataset of negative examples. While this ratio favors the classification of the positive class, it does not take into account that in a real-life scenario of a microbial fermentation the peptide that is generated that does not belong to the positive class is not random but is more dependent on the substrate utilized. For example, if the substrate to be used is from a protein matrix derived from grains, the amount of celiac disease peptides would be much higher than the rest of the other classes.For this reason, in the case of microbial fermentation, the choice of the positive/negative dataset plays an important role in order to get closer to the real-life scenario. CICERON adopts a distinct strategy since it considers peptides from other functional classes as negative datasets, thus enhancing its ability to discern both positive and negative sample characteristics.

An additional distinction lies in the peptide length criteria, where each class has predefined minimum and maximum sequence length constraints (Table 2), thereby restricting the classification to peptides within those specific ranges for SOTA classifiers. If enzymatic proteolysis or physicochemical proteolysis are also compared, the length of the peptides obtained is different than in the case of microbial fermentation, where only certain microbial enzymes are present and thus the digested proteins produce longer peptides. This highlights the need to have separate datasets for each proposed proteolysis method in order to obtain more accurate predictions similar to a real experiment.

Upon a comprehensive review of all SOTA classifiers, it emerges there is not a common encoding or classification methodology shared among these models. If the objective is to obtain a specific class of peptide, i.e. antimicrobial peptides are needed from fermentation, a class-specific classifier can in principle be more suited to the case. However, it cannot be excluded that a function-specific classifier is not able to see if the peptide has more than one function. In one extreme case, if the peptide is antihypertensive but at the same time it is associated with celiac disease, it could cause problems for a patient suffering from the disease. A preliminary screening with CICERON could help identify these cases very early in the screening process, eliminating the need for costly and time-consuming tests.

## Conclusions

To the best of the authors’ knowledge, this is the first systematic benchmark specifically focusing on the functional classification of BPs by utilizing a comprehensive dataset of peptides derived from food fermentation. Taking into account the importance of fermentation for the production of bioactive molecules, the generation of classifiers for the identification of functions will facilitate the identification of new BP. In this study, we successfully established new machine learning classifiers for previously unaddressed functional classes, including opioids, cardiovascular peptides, and immunomodulatory peptides. Furthermore, classifiers for antidiabetic, celiac disease and antioxidative peptides demonstrated superior performance on a test close to a real-case scenario compared to previously developed models. Nevertheless, certain classes, such as antidiabetic and immunomodulatory peptides, still present challenges in terms of classification performance, highlighting the importance of expanding the number of sequences in these groups. Our comprehensive analysis, encompassing diverse encoding techniques and classification methodologies, unveiled the unique requirements of each functional class for accurate functional identification. It is evident that there is no one-size-fits-all solution for all classes. Moreover, the examination of amino acid composition revealed distinctive preferences within certain groups, enabling discrimination based on specific sequence motifs. While these attributes do not exclusively determine functional group affiliation, they can significantly enhance the classification efficacy of the classifiers. From the present study, it clearly emerges that conventional methods such as LR can be effectively employed in classification tasks, and that more advanced algorithms such as NN do not necessarily provide better results. Notably, the use of language models, such as ProtBERT, proves to be a straightforward method for functional classification, although the performance can be lower in comparison to algorithms specifically crafted for the problem. These pre-trained models require minimal optimization, except for the selection of the appropriate output layer that should be tailored to the specific problem. In the context of a microbial fermentation experiment, where the goal is to characterize the full spectrum of peptides present within the substrate, a versatile tool like CICERON proves to be more efficient than SOTA classifiers in identifying multiple functions associated with one single peptide, making it a valuable tool for exploratory investigations into peptide functions within complex biological systems.

## Data and code availability

The database used in this study, the scripts to reproduce the analyses, and the best model for each functional class can be found at https://github.com/BizzoTL/CICERON/.

## Authors contributions

Conceptualization, E.B, G.Z and S.C; methodology, E.B and G.Z; software, E.B; formal analysis, E.B; investigation, E.B; data curation, E.B; writing—original draft, E.B; writing—review and editing, E.B, G.Z, L.T, P.F, R.D.C and S.C.; supervision, S.C. and L.T.; fundings P.F, R.D.C, S.C.; all the authors read and approved the final manuscript.

## Author Biographies

**Edoardo Bizzotto** is a PhD student at the Department of Biology at the University of Padova. His research interests include machine learning with a focus on bioinformatics and microbial metagenomics.

**Guido Zampieri** is a Junior Researcher at the Department of Biology at the University of Padova. His research interests include the application of machine learning and genome-scale modeling in the context of microbial genomics and metagenomics.

**Laura Treu** is Senior Researcher at the Department of Biology at the University of Padova. Her research activity is focused on the use of microorganisms in the context of anaerobic digestion and circular economy.

**Pasquale Filannino** is Associate Professor at the Department of Soil, Plant and Food Science at the University of Bari Aldo Moro. His research activity is focused on microbial systems biotechnology, fermented and functional foods, and microbial physiology.

**Raffaella Di Cagno** is Full Professor at the Faculty of Agricultural, Environmental and Food Sciences at the University of Bolzano. Her research is focused on genomics and proteomics of lactic acid bacteria (LAB), the effect, and metabolism of LAB and related food biotechnology applications, and the effect of the diet on gut microbiome and human health.

**Stefano Campanaro** is Associate Professor at the Department of Biology at the University of Padova. His research activity is mainly focused on bioinformatics in the context of genomics and metagenomics and he has worked with microorganisms fermentation.

## References

1. Bhandari D, Rafiq S, Gat Y, et al. A Review on Bioactive Peptides: Physiological Functions, Bioavailability and Safety. Int. J. Pept. Res. Ther. 2020; 26:139–150

2. Sánchez A, Vázquez A. Bioactive peptides: A review. Food Qual. Saf. 2017; 1:29–46

3. Perlikowska R, Silva J, Alves C, et al. The Therapeutic Potential of Naturally Occurring Peptides in Counteracting SH-SY5Y Cells Injury. Int. J. Mol. Sci. 2022; 23:11778

4. da Costa JP, Cova M, Ferreira R, et al. Antimicrobial peptides: an alternative for innovative medicines? Appl. Microbiol. Biotechnol. 2015; 99:2023–2040

5. Da Silva J, Leal EC, Carvalho E. Bioactive Antimicrobial Peptides as Therapeutic Agents for Infected Diabetic Foot Ulcers. Biomolecules 2021; 11:1894

6. Haney EF, Mansour SC, Hancock REW. Antimicrobial Peptides: An Introduction. Antimicrob. Pept. Methods Protoc. 2017; 3–22

7. Minkiewicz P, Iwaniak A, Darewicz M. BIOPEP-UWM Database of Bioactive Peptides: Current Opportunities. Int. J. Mol. Sci. 2019; 20:5978

8. Cheung RCF, Ng TB, Wong JH. Marine Peptides: Bioactivities and Applications. Mar. Drugs 2015; 13:4006–4043

9. Chakrabarti S, Guha S, Majumder K. Food-Derived Bioactive Peptides in Human Health: Challenges and Opportunities. Nutrients 2018; 10:1738

10. Kitts DD, Weiler K. Bioactive Proteins and Peptides from Food Sources. Applications of Bioprocesses used in Isolation and Recovery. Curr. Pharm. Des. 9:1309–1323

11. Costa EM, Oliveira AS, Silva S, et al. Spent Yeast Waste Streams as a Sustainable Source of Bioactive Peptides for Skin Applications. Int. J. Mol. Sci. 2023; 24:2253

12. Harnedy PA, FitzGerald RJ. Bioactive peptides from marine processing waste and shellfish: A review. J. Funct. Foods 2012; 4:6–24

13. Akbarian M, Khani A, Eghbalpour S, et al. Bioactive Peptides: Synthesis, Sources, Applications, and Proposed Mechanisms of Action. Int. J. Mol. Sci. 2022; 23:1445

14. Kadam SU, Tiwari BK, Álvarez C, et al. Ultrasound applications for the extraction, identification and delivery of food proteins and bioactive peptides. Trends Food Sci. Technol. 2015; 46:60–67

15. Cruz-Casas DE, Aguilar CN, Ascacio-Valdés JA, et al. Enzymatic hydrolysis and microbial fermentation: The most favorable biotechnological methods for the release of bioactive peptides. Food Chem. Mol. Sci. 2021; 3:100047

16. Najafian L, Babji AS. Production of bioactive peptides using enzymatic hydrolysis and identification antioxidative peptides from patin (Pangasius sutchi) sarcoplasmic protein hydolysate. J. Funct. Foods 2014; 9:280–289

17. Sharma P, Sosalagere C, Kehinde BA, et al. Chapter 15 - Bioactive peptides production using microbial resources. Microb. Resour. Technol. Sustain. Dev. 2022; 299–317

18. Raveschot C, Cudennec B, Coutte F, et al. Production of Bioactive Peptides by Lactobacillus Species: From Gene to Application. Front. Microbiol. 2018; 9:

19. Du Z, Ding X, Xu Y, et al. UniDL4BioPep: a universal deep learning architecture for binary classification in peptide bioactivity. Brief. Bioinform. 2023; bbad135

20. Fang Y, Xu F, Wei L, et al. AFP-MFL: accurate identification of antifungal peptides using multi-view feature learning. Brief. Bioinform. 2023; 24:bbac606

21. Sharma R, Shrivastava S, Kumar Singh S, et al. Deep-AFPpred: identifying novel antifungal peptides using pretrained embeddings from seq2vec with 1DCNN-BiLSTM. Brief. Bioinform. 2022; 23:bbab422

22. Sharma R, Shrivastava S, Kumar Singh S, et al. Deep-ABPpred: identifying antibacterial peptides in protein sequences using bidirectional LSTM with word2vec. Brief. Bioinform. 2021; 22:bbab065

23. Tang W, Dai R, Yan W, et al. Identifying multi-functional bioactive peptide functions using multi-label deep learning. Brief. Bioinform. 2022; 23:bbab414

24. Zhang Y, Lin J, Zhao L, et al. A novel antibacterial peptide recognition algorithm based on BERT. Brief. Bioinform. 2021; 22:bbab200

25. Chaudhary A, Bhalla S, Patiyal S, et al. FermFooDb: A database of bioactive peptides derived from fermented foods. Heliyon 2021; 7:e06668

26. PepCalc.com - Peptide property calculator. 2015;

27. Dai R, Zhang W, Tang W, et al. BBPpred: Sequence-Based Prediction of Blood-Brain Barrier Peptides with Feature Representation Learning and Logistic Regression. J. Chem. Inf. Model. 2021; 61:525–534

28. Meng C, Jin S, Wang L, et al. AOPs-SVM: A Sequence-Based Classifier of Antioxidant Proteins Using a Support Vector Machine. Front. Bioeng. Biotechnol. 2019; 7:

29. Zhao D, Teng Z, Li Y, et al. iAIPs: Identifying Anti-Inflammatory Peptides Using Random Forest. Front. Genet. 2021; 12:

30. Chen J, Cheong HH, Siu SWI. xDeep-AcPEP: Deep Learning Method for Anticancer Peptide Activity Prediction Based on Convolutional Neural Network and Multitask Learning. J. Chem. Inf. Model. 2021; 61:3789–3803

31. Lei Y, Li S, Liu Z, et al. A deep-learning framework for multi-level peptide–protein interaction prediction. Nat. Commun. 2021; 12:5465

32. Spänig S, Mohsen S, Hattab G, et al. A large-scale comparative study on peptide encodings for biomedical classification. NAR Genomics Bioinforma. 2021; 3:lqab039

33. Spänig S, Heider D. Encodings and models for antimicrobial peptide classification for multi-resistant pathogens. BioData Min. 2019; 12:7

34. Sidorczuk K, Gagat P, Pietluch F, et al. Benchmarks in antimicrobial peptide prediction are biased due to the selection of negative data. Brief. Bioinform. 2022; 23:bbac343

35. Nielsen SD, Beverly RL, Qu Y, et al. Milk bioactive peptide database: A comprehensive database of milk protein-derived bioactive peptides and novel visualization. Food Chem. 2017; 232:673–682

36. Li Q, Zhang C, Chen H, et al. BioPepDB: an integrated data platform for food-derived bioactive peptides. Int. J. Food Sci. Nutr. 2018; 69:963–968

37. Vens C, Rosso M-N, Danchin EGJ. Identifying discriminative classification-based motifs in biological sequences. Bioinformatics 2011; 27:1231–1238

38. Maetschke S, Towsey M, Boden M. BLOMAP: An Encoding Of Amino Acids Which Improves Signal Peptide Cleavage Site Prediction. Proc. 3rd Asia-Pac. Bioinforma. Conf. Adv. Bioinforma. Comput. Biol. 2005; 141–150

39. Brandes N, Ofer D, Peleg Y, et al. ProteinBERT: a universal deep-learning model of protein sequence and function. Bioinformatics 2022; 38:2102–2110

40. Maaten L van der, Hinton G. Visualizing Data using t-SNE. J. Mach. Learn. Res. 2008; 9:2579–2605

41. Pedregosa F, Varoquaux G, Gramfort A, et al. Scikit-learn: Machine Learning in Python. J. Mach. Learn. Res. 2011; 12:2825–2830

42. Chollet, F. & others. Keras: Deep Learning for humans. 2015;

43. Wolf T, Debut L, Sanh V, et al. HuggingFace’s Transformers: State-of-the-art Natural Language Processing. 2020;

44. De Rainville F-M, Fortin F-A, Gardner M-A, et al. DEAP: a python framework for evolutionary algorithms. Proc. 14th Annu. Conf. Companion Genet. Evol. Comput. 2012; 85–92

45. Tomer R, Patiyal S, Dhall A, et al. Prediction of celiac disease associated epitopes and motifs in a protein. Front. Immunol. 2023; 14:1056101

46. Chen X, Huang J, He B. AntiDMPpred: a web service for identifying anti-diabetic peptides. PeerJ 2022; 10:e13581

47. Qin D, Jiao L, Wang R, et al. Prediction of antioxidant peptides using a quantitative structure−activity relationship predictor (AnOxPP) based on bidirectional long short-term memory neural network and interpretable amino acid descriptors. Comput. Biol. Med. 2023; 154:106591

48. Akbar S, Mohamed HG, Ali H, et al. Identifying Neuropeptides via Evolutionary and Sequential Based Multi-Perspective Descriptors by Incorporation With Ensemble Classification Strategy. IEEE Access 2023; 11:49024–49034

49. Cao Q, Ge C, Wang X, et al. Designing antimicrobial peptides using deep learning and molecular dynamic simulations. Brief. Bioinform. 2023; 24:bbad058

50. Lertampaiporn S, Hongsthong A, Wattanapornprom W, et al. Ensemble-AHTPpred: A Robust Ensemble Machine Learning Model Integrated With a New Composite Feature for Identifying Antihypertensive Peptides. Front. Genet. 2022; 13:

