## Supplementary Materials for "Classification of bioactive peptides: a comparative analysis of models and encodings"

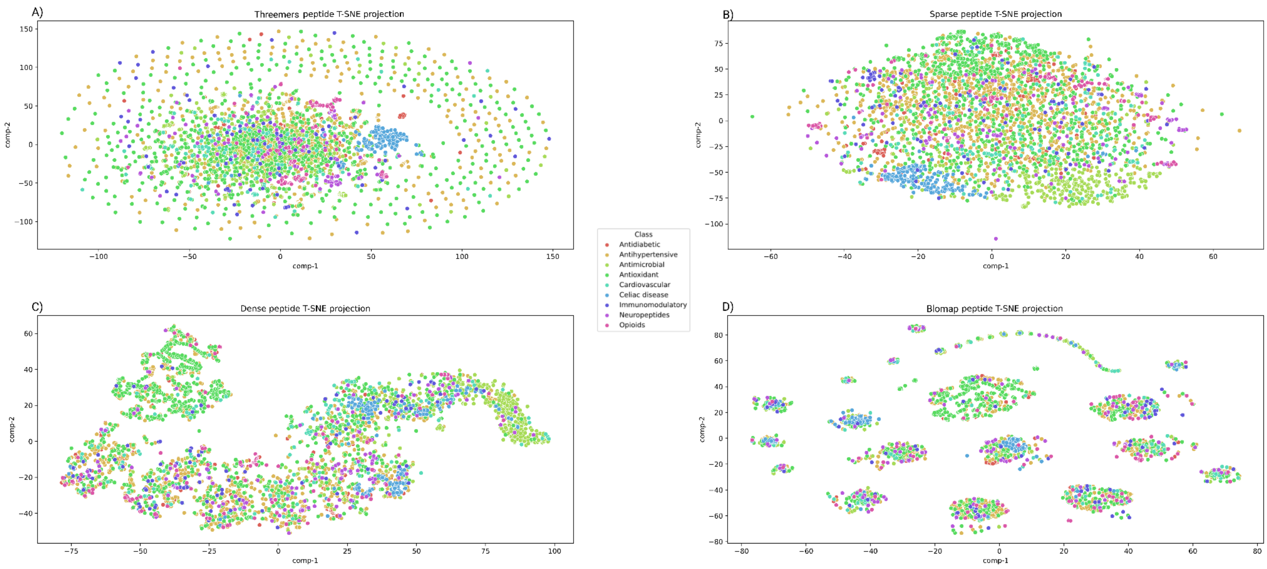


**Supplementary Figure 1.** T-SNE representation of the four different peptide encodings: A) Threemers encoding, B) Sparse encoding, C) Dense encoding, D) Blomap encoding.

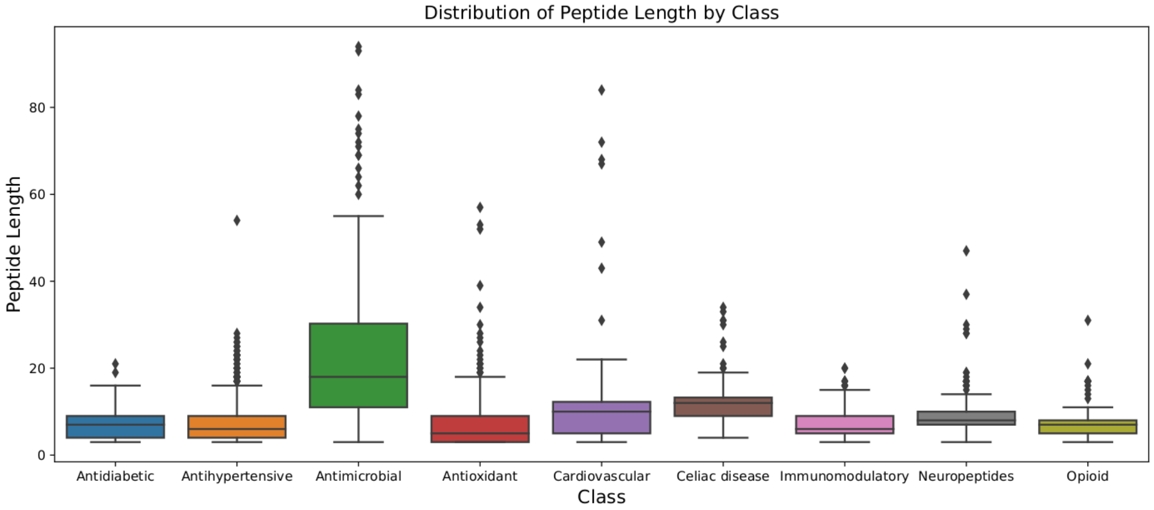


**Supplementary Figure 2.** Distribution of the average peptide length for each functional class.

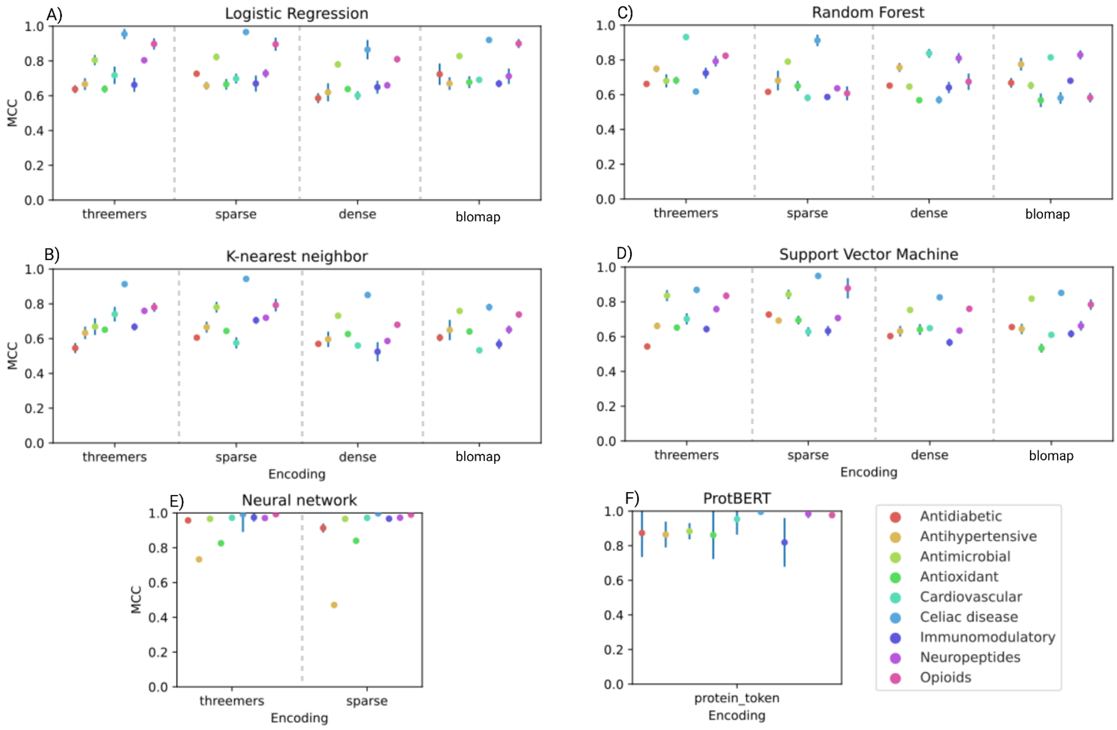


**Supplementary Figure 3.** AUROC value on the test set for every single classifier and encoding method. The dot size corresponds to a value of 0.03 AUROC. Standard deviation values lower than 0.03 are not displayed. Each plot describes the performance of a different machine learning method: A) Logistic Regression, B) K-Nearest Neighbor, C) Random Forest, D) Support Vector Machine, E) Neural Network, F) ProtBERT.

**Supplementary Table 1**

Training and test metrics for each classifier and encoding method used in this work. The best parameters found in the optimization step are also listed for LR, RF, SVM and KNN. For NN the best individual in all the generations is reported.

**Supplementary Table 2**

Amino acids, dipeptides, tripeptides motifs and the corresponding p-values obtained with binomial tests. MERCI motifs are also reported.
